# Anti-CD3 microporous annealed particle hydrogel protects stem cell derived beta cells from autoreactive T cells

**DOI:** 10.1101/2025.11.11.687837

**Authors:** Adrienne E. Widener, Cameron T. Manson, Jessie M. Barra, Christopher P. Spencer, Amanda D. Grodman, Andrew M. Ladd, Holger A. Russ, Edward A. Phelps

## Abstract

Type 1 diabetes (T1D) results from autoimmune destruction of pancreatic beta cells, leaving patients dependent on exogenous insulin and at risk of severe hypoglycemic episodes. Stem cell-derived beta-like cells (sBCs) offer a promising approach for beta cell replacement therapy, but clinical translation is limited by immune-mediated rejection, recurrent autoimmunity, and inhospitable transplantation sites. Biomaterials have been investigated to provide localized immune-isolation and immunomodulation, but foreign body responses and rapid depletion of therapeutic agents remain as obstacles to clinical translation. Here, we present a microporous annealed particle (MAP) hydrogel functionalized with an anti-CD3 monoclonal antibody (αCD3) to provide a localized immunomodulatory microenvironment for beta cell replacement therapy. MAP hydrogels consisting of guest-host interlinked polyethylene glycol-maleimide (PEG-MAL) microgels supported rapid vascularization, minimal foreign body response, and engraftment of syngeneic islets in mice. αCD3 MAP hydrogel halted T cell migration *in vitro* and protected transplanted sBCs from immune-mediated destruction by HLA-matched diabetogenic T cells *in vivo*. Subcutaneous αCD3 functionalized MAP hydrogel also protected the endogenous islets in the pancreas, demonstrating potential for systemic immune modulation. These findings establish αCD3 MAP hydrogels as a promising strategy for localized immune modulation in cell replacement therapy.

## Introduction

Type 1 diabetes (T1D) is a chronic disease characterized by the immune-mediated destruction of insulin-producing pancreatic beta cells, leaving patients dependent on exogenous insulin therapy. Despite advances in diabetes management technology, patients remain at risk for severe hypoglycemic episodes and secondary complications of chronic hyperglycemia. [1] Beta cell replacement therapies using the Edmonton protocol, which enabled the isolation and transplantation of human islets from cadaveric donors, demonstrated insulin independence in early clinical trials and sparked renewed interest in beta cell replacement as a therapeutic strategy. [2, 3] However, clinical application is severely limited due to limited donor tissue availability, highlighting the need for an alternative source of insulin-producing beta cells. The generation of stem cell-derived beta-like cells (sBCs) holds tremendous promise as a personalized and scalable source of functional, insulin-producing cells for beta cell replacement therapy. Results from initial clinical trials using sBCs have reported normalized blood sugar levels in addition to insulin independence in 83% of patients 90 days post-transplant. [4] Graft rejection of beta cell replacement therapies by allorecognition and recurrence of autoimmunity is currently prevented by broad-spectrum immunosuppressive agents. However, long-term use of immunosuppressants increases risk for opportunistic infections, malignancies, and drug-related toxicities, particularly to the kidneys. [5] In addition, the use of the liver as an engraftment site can lead to complications of thrombosis, instant blood-mediated inflammatory responses, and amyloidosis, ultimately leading to graft loss. Alternative sites such as the omentum and subcutaneous space hold potential but are limited by delayed vascularization, resulting in ischemia of the beta cell graft, and differences in the extracellular microenvironment from the native pancreas lead to beta cell loss by anoikis. [6]

Biomaterials have long been used for encapsulation of cellular therapies and for localized delivery of immune modulation. [7, 8] However, biomaterials face challenges posed by the foreign body response and rapid depletion of loaded immunomodulatory agents. [7, 9] Fine-tuning of biomaterial properties such as porosity, stiffness, and surface chemistry has been shown to reduce the foreign body response and promote graft survival in multiple transplant sites. [10] Microporous annealed particle (MAP) hydrogels are a class of biomaterial formed by compacting small microgels into a larger interconnected scaffold through secondary cross-links or an annealing procedure. MAP hydrogels are a subclass of the broader category of granular hydrogels, characterized by the interparticle annealing that increases material strength and cohesiveness. [11] MAP hydrogels have shown potential to improve engraftment and survival of transplanted beta cells by promoting rapid vascularization and efficient nutrient exchange, and can be engineered to present tethered immunomodulatory proteins. [12, 13] This capability allows for localized and sustained engagement of immune cell subsets and supports long-term graft survival. Together, these features position MAP hydrogels as a promising next-generation platform for cell replacement therapies in T1D.

In this study, we developed an immunomodulatory guest-host interlinked PEG-based microgel platform conjugated with an anti-CD3 monoclonal antibody (αCD3) to prevent autoimmune rejection of transplanted sBCs in a subcutaneous site. Previously, we established a MAP hydrogel interlinked with guest-host molecules (guest-host MAP) as an injectable microenvironment for the delivery of islets and to enable rapid cell infiltration. [14–16] Here we demonstrate minimal foreign body response to this material and the development of vasculature in the subcutaneous space. We conjugated αCD3 to the surfaces of guest-host MAP hydrogel (αCD3 MAP) to attenuate autoreactive T cell responses in a human-HLA mouse model of T1D. Teplizumab is a humanized monoclonal antibody therapy for T1D prevention that binds specifically to the CD3 complex on T cells, leading to their functional inactivation and deletion. Teplizumab, a modified form of αCD3 with reduced Fc receptor binding, is the first FDA-approved drug to delay the onset of clinical T1D in high-risk individuals. [17] In its clinical trials (TrialNet 10), a single course of Teplizumab significantly slowed progression to clinical T1D in high-risk individuals who had at least two autoantibodies. [18] Soluble αCD3 has also shown efficacy in prolonging graft survival in allogeneic transplantation models. [19] Here, we hypothesize that the use of biomaterial-tethered αCD3 would localize the immunosuppressive capacity of αCD3 to the beta cell graft site. To investigate the protection of transplanted human sBCs by αCD3 MAP in a reproducible manner, we established an HLA-matched humanized T1D model using NOD-cMHCI^-^-A2, which lacks endogenous MHC class I genes and expresses a human HLA-A2 transgene, in combination with NSG-HLA-A2/HHD mice, which carry the same transgene with the addition of a SCID mutation, that renders them immunodeficient. CD8^+^ T cells isolated from diabetic NOD-cMHCI^-^-A2 mice can directly interact and engage with HLA-A2 expressed on human beta cells. [20] αCD3 MAP arrested the migration of activated T cells *in vitro*. We adoptively transferred CD3^+^ T cells from diabetic NOD-cMHCI^-^-A2 mice to NSG-HLA-A2/HHD mice to create a controlled induction of autoimmune diabetes. sBCs transplanted with αCD3 MAP in immunodeficient mice were resistant to immune challenge by adoptive transfer of diabetogenic T cells. Intriguingly, subcutaneous αCD3 MAP conferred remote protection against immune-mediated destruction of islets in the pancreas. The approach demonstrates a localized immune modulation approach that has the potential to improve engraftment and survival of beta cell replacement therapies without systemic immune suppression.

## Results

### Guest-host MAP hydrogels have shear-reversible cross-links

Here, we developed MAP hydrogels as a supportive biomaterial platform to enhance the survival and engraftment of transplanted sBCs. Previously, we adapted our islet-compatible polyethylene glycol maleimide (PEG-MAL) hydrogel technology [21–23] for the formation of guest-host MAP microgels through batch emulsion. [14–16] In brief, four-arm PEG-MAL macromer is pre-functionalized with adamantane-mono-thiol (guest) and β-cyclodextrin-mono-thiol (host) before the addition of a four-arm PEG-thiol (PEG-SH) crosslinker. We then generate a water-in-oil emulsion by vortexing, allow the emulsion droplets to complete the cross-linking reaction, and collect and wash the formed microgels by centrifugation. The microgels, exhibiting both guest and host molecules on each particle, are inter-linked when compacted by the guest-host interactions of the adamantane and β-cyclodextrin groups present on the surface (**Figure 1A**). The guest-host interactions are also present in the bulk of the hydrogel network and contribute to the material properties. [14–16] The emulsion method generated PEG microgels of average diameter 19.4 μm and polydispersity index (PDI) of 0.29, and was reproducible across multiple batches (**Figure 1B, C**).

**Figure 1:**
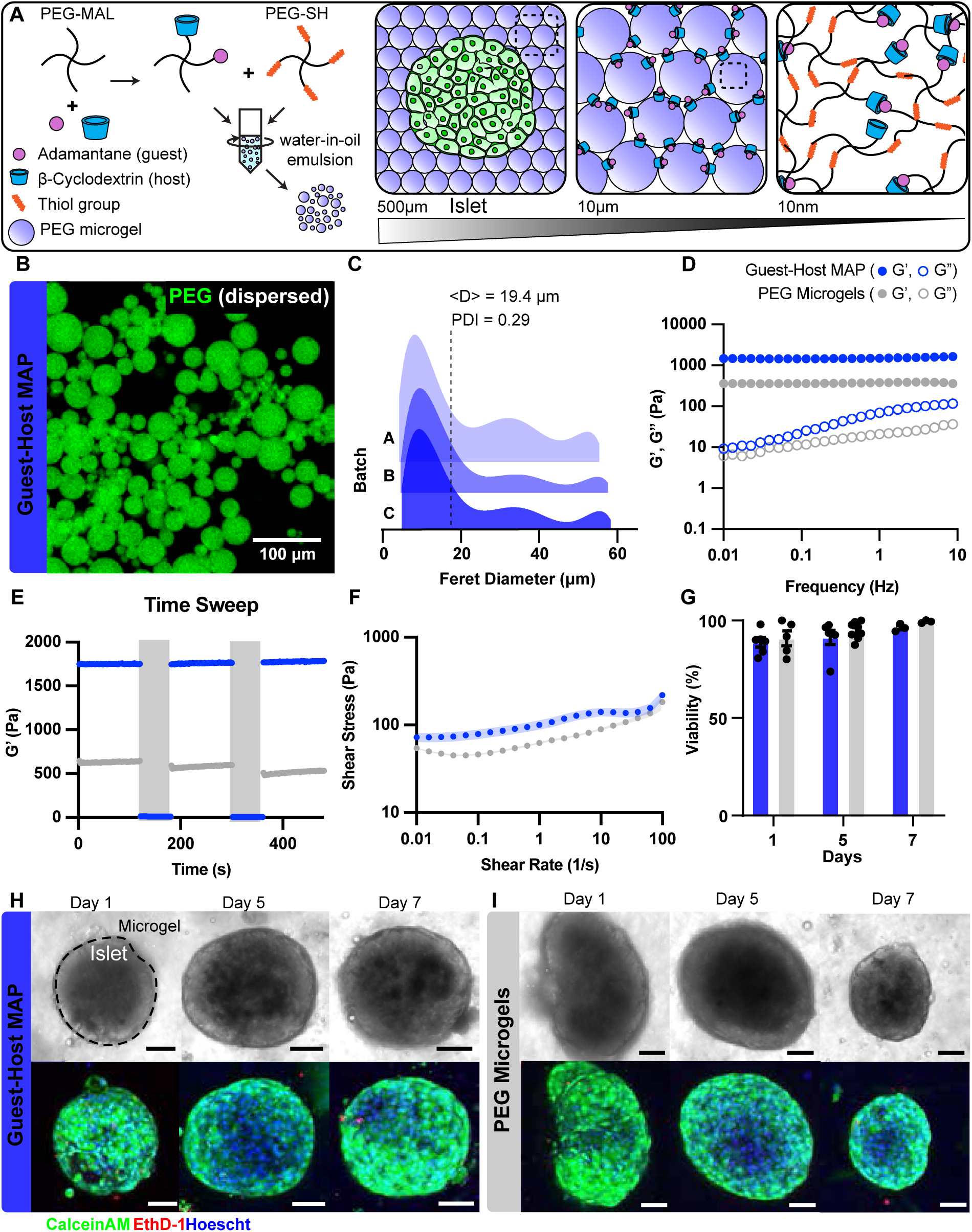
Guest-host MAP hydrogels have shear-reversible cross-links. (A) Schematic of the approach to generate guest-host MAP. Four-arm PEG-MAL was reacted with adamantane-thiol (guest) and mono-thiol β-cyclodextrin (host). The functionalized PEG macromer was mixed with PEG-SH and vortexed to form a water-in-oil emulsion to create microgels. Microgels are close-packed to form a porous scaffold with embedded cells or pancreatic islets (left). The guest-host MAP (middle) is made up of many interlinked microspheres packed together, creating interstitial spaces allowing for porosity and guest-host crosslinks interacting between individual microspheres. The structure of the guest-host MAP (right) is constructed of both physical guest-host and chemical maleimide-thiol crosslinks. (B) Confocal image of guest-host MAP. PEG was labeled with Alexa Fluor 647 C5-maleimide during microgel fabrication. Microgels are dispersed for visualization and size measurements. Scale bar is 100 μm. (C) Feret diameter of microgels made by batch emulsion. The average diameter for all batches is 19.4 μm, and the polydispersity index (PDI) is 0.29. (D) Oscillatory frequency sweep from 10 to 0.01 Hz at 1% amplitude for guest-host MAP (blue) and unfunctionalized PEG microgels (grey). Data is represented by means of technical replicates (n = 3) (E) Unidirectional shear rate sweep from 100 1/s to 0.01 1/s. Data is represented by mean ± SD of technical replicates (n = 3) (F) Cyclic strain alternating between low (1%) for 120s and high (500%) strain for 60 s. Data is represented by means of technical replicates (n = 3) (G). Quantified viability of human islets cultured in guest-host MAP and PEG microgels over 7 days. Data represents mean ± SEM. Data was analyzed by two-way ANOVA with multiple comparisons (n = 3-5). Transmitted light (top) and fluorescent (bottom) images of mouse islets cultured in (H) guest-host MAP and (I) unfunctionalized PEG microgels over days 1, 5, and 7. Islets labeled with a dotted line indicating separation of microgels and embedded islets. Islets were stained with CalceinAM (Live), EthD-1 (dead), and Hoescht (nuclei). Scale bar is 50μm.

To assess the rheological properties of guest-host MAP, we performed a low amplitude (1%) oscillatory frequency sweep from 10 to 0.01 Hz and found that macroscopic scaffolds composed of PEG-MAL microgels, both with and without guest-host, displayed viscoelastic behavior over the entire frequency range (**Figure 1D**). The guest-host MAP had a significantly higher storage modulus (1500 Pa) than unfunctionalized PEG microgels without guest-host (390 Pa) over the entire frequency range. We next showed that guest-host MAP hydrogel (blue) demonstrated full recovery of the storage modulus following repeated high-strain deformation (**Figure 1E**) while maintaining low yield stresses (**Figure 1F**), indicating shear-reversible thinning and self-healing properties that make the material suitable for injectable delivery and mechanical resilience in a transplant setting. In contrast, PEG microgels (grey) without guest-host progressively lost structural integrity upon repeated exposure to high strains, indicating a lack of self-healing capacity (**Figure 1E, F**). We next investigated the cytocompatibility of the guest-host MAP by culturing C57BL/6J mouse islets embedded in the interstitial spaces between microgels. Islet viability was excellent for both guest-host MAP and PEG microgels, with no decline observed over seven days in culture (**Figure 1G**). In both microgel types, islets maintained a rounded morphology with minimal cell shedding throughout the culture period (**Figure 1H, I**). These results indicate that the addition of guest-host interactions had no effect on the viability of embedded islets within the hydrogel.

### Guest-host MAP promotes subcutaneous vascularization

We hypothesized that the microporous architecture and guest-host interactions of guest-host MAP would minimize the foreign body response and promote vascularization in the subcutaneous space. It has been previously demonstrated that cell-scale porosity with appropriate mechanical stability promotes a healing response. [24] To assess the ability of our microgels to support islets transplanted in the subcutaneous space, guest-host MAP and PEG microgels were injected subcutaneously into C57BL/6 mice without islet cargo. Implants were retrieved for histological analysis at day 31. Both microgel types exhibited low fibrotic encapsulation (capsule thickness PEG microgels ∼ 100 µm, guest-host MAP ∼ 31 µm) and showed evidence of integration with the surrounding host tissue via cellular infiltration and tissue remodeling (**Figure 2A, B**). There was a marked difference in cellular infiltration and vascular integration between guest-host MAP and PEG microgels. In guest-host MAP implants, infiltrating cells were organized into elongated, vessel-like structures spanning the scaffold and were positive for basement membrane markers (**Figure 2C**), while such structures were absent from PEG microgels without guest-host. Cellular infiltration into the guest-host MAP hydrogel extended approximately 19.2 µm from the surface, whereas cells within the PEG microgels exhibited a more uniform and dense distribution, with an average infiltration of 59.4 µm (**Figure 2D, E**). To confirm vascular integration, we stained for CD31 and found a greater density of vessels and a significant increase in CD31^+^ area within guest-host MAP hydrogels (**Figure 2F, G**). These results indicate that guest-host MAP facilitates blood vessel formation and tissue remodeling, which are critical for the survival and function of transplanted beta cells.

**Figure 2:**
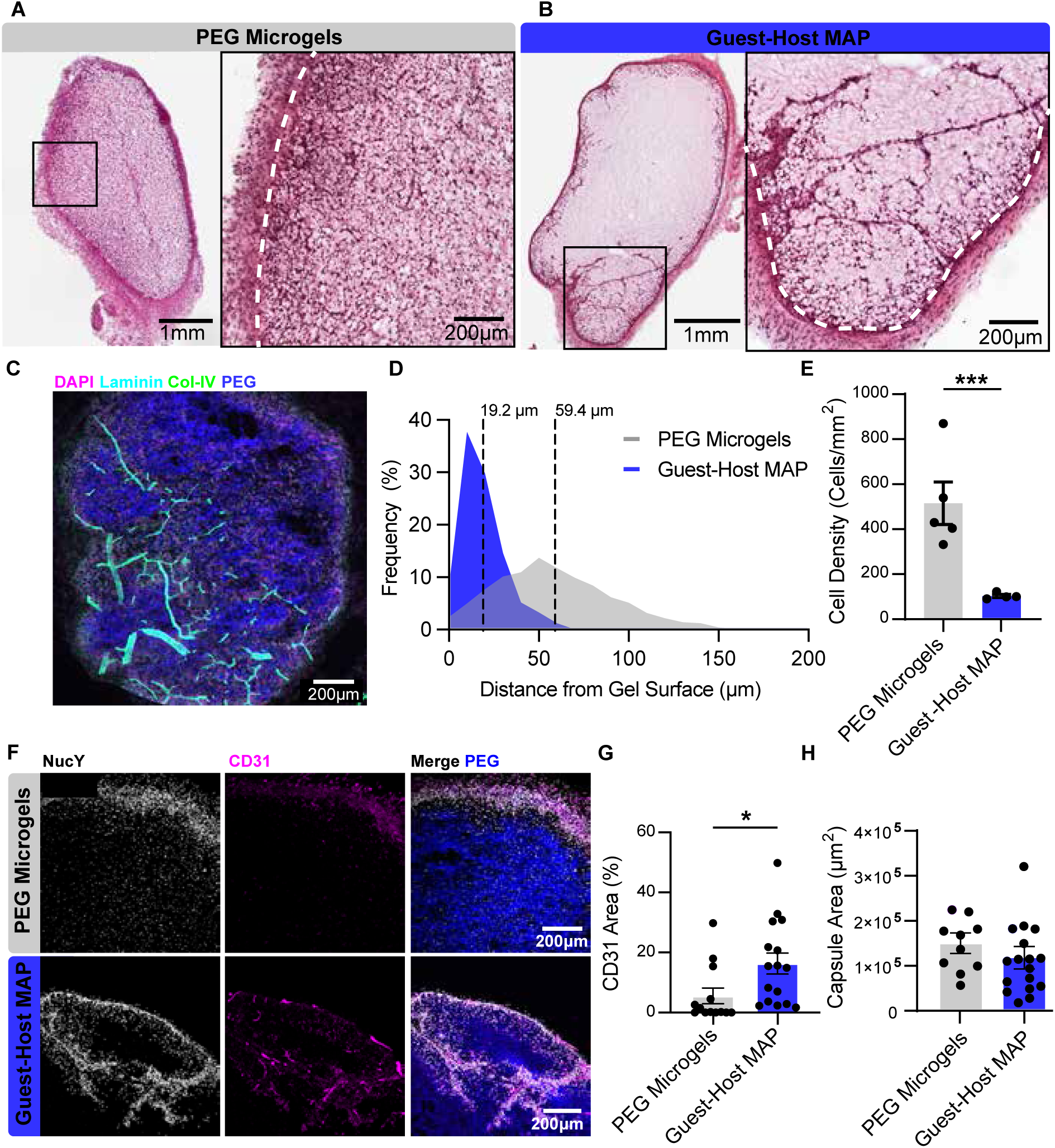
Guest-host MAP promotes subcutaneous vascularization. H&E staining of (A) PEG microgel and (B) guest-host MAP implants (left) and magnification of the tissue interface (right) at 31 days post-transplantation. The white dotted line represents the border of the gel. (C) Confocal image of guest-host MAP implant with immunofluorescence staining for laminin and collagen IV and co-stained with DAPI (nuclei). PEG was labeled with Alexa Fluor 647 C5-maleimide during microgel fabrication. (D) Histogram of the distance of infiltrating cell nuclei from the gel surface for guest-host MAP and PEG microgels. The average distance of nuclei infiltrating into the implant from the gel surface is 19.2 μm for guest-host MAP and 59.4 μm for PEG microgels. (E) Quantification of infiltrating cell density expressed as the number of nuclei per unit area of the implant. (F) Vascularization of guest-host MAP and PEG microgels in the subcutaneous space after 31 days. NucY (nuclear yellow stain), CD31 (endothelial cells), and PEG (blue). (G) Percent CD31^+^ area per unit area of implant for guest-host MAP and PEG microgels. (H) Area of the fibrous capsule surrounding the guest-host MAP and PEG microgel implants. Each dot represents a single image. Data represented by mean ± SEM and analyzed via nested t-test of multiple sections, each of four independent biological replicates (N=4 mice, n = 3-4 sections per mouse). *p < 0.05, ** p < 0.01, *** p < 0.001.

### Subcutaneous engraftment of syngeneic islet transplants in guest-host MAP

To evaluate the functional capacity of islets transplanted in guest-host MAP in hyperglycemic conditions, we used a streptozotocin (STZ)-induced model of diabetes in C57BL/6 mice. Syngeneic mouse islets were encapsulated within guest-host MAP hydrogels and injected subcutaneously. Blood glucose was monitored longitudinally to assess graft function and restoration of normoglycemia (defined as two subsequent readings < 300 mg/dl) (**Figure 3A**). Mice receiving islets within guest-host MAP hydrogels achieved non-fasting blood glucose below 300 mg/dL by six weeks and below 250 mg/dL by eight weeks (**Figure 3B**), bare islets transplanted subcutaneously without a biomaterial failed within one week (**Figure 3B**). Intraperitoneal glucose tolerance tests revealed moderate glycemic control at six weeks and no significant difference between guest-host MAP and non-diabetic controls by eight weeks (**Figure 3C-E**). Grafts were explanted after 70 days for histological analysis, revealing vascularized islets within the guest-host MAP with extensive intra-islet CD31^+^ area (**Figure 3F**). These findings demonstrate that guest-host MAP provides a supportive subcutaneous microenvironment that promotes islet engraftment, vascular integration, and functional recovery to normoglycemic levels.

**Figure 3.**
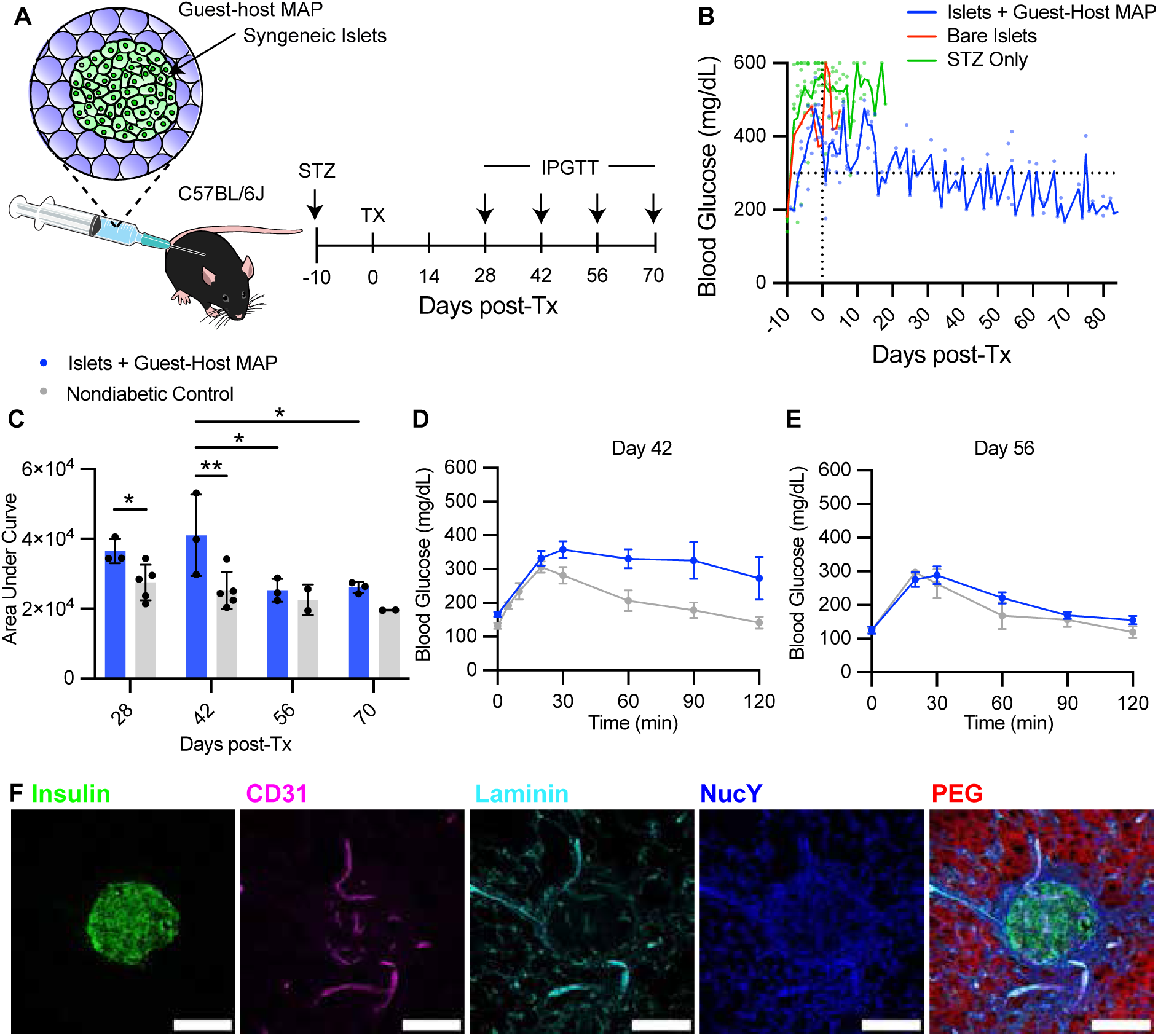
Syngeneic islets co-delivered with MAP-PEG engraft subcutaneously in diabetic mice. (A) Schematic of the experimental overview. Four hundred syngeneic islets were injected in 200 µl guest-host MAP and injected into the dorsal subcutaneous space of STZ-induced diabetic C57BL/6J mice. (Right) Experimental timeline relative to the day of transplant, with STZ administration on day-10 and IPGTTs on days 28, 42, 56, and 70 post-transplantation. (B) Non-fasting blood glucose measurements for mice receiving islets + guest-host MAP (n=3) or bare islets (n=1). After day 0, each data point represents the 7-day mean non-fasting blood glucose. Error bars = mean ± SEM. (C) Area under the curve measurements for individual IPGTTs conducted at days 28, 42, 56, and 70 post-transplantation. Transplant recipients (blue) are compared to nondiabetic controls (grey). Data analyzed by two-way ANOVA with Tukey’s post-hoc multiple comparisons. Mean ± SEM, *p < 0.05, ** p < 0.01 (D-E) IPGTT curves recorded at days 42 and 56 post-transplantation. Mean ± SEM. Islets + guest-host MAP (n=3), nondiabetic control (n=2). (F) Representative confocal images of immunostaining of an islet engrafted in guest-host MAP after graft removal. Sections were stained for insulin, CD31, and laminin. Nuclear yellow was used as a nuclear counterstain. PEG was labeled with Alexa Fluor 647 C5-maleimide during microgel fabrication. Scale bar = 100 µm.

### Synthesis of αCD3 guest-host microgels that arrest T cell migration

Next, we evaluated the effect of functionalizing guest-host MAP with αCD3 to arrest T cell migration. To functionalize guest-host MAP with αCD3, we generated amine-reactive microgels. To do this, we pre-reacted the four-arm PEG-SH crosslinker with Sulfo-SMCC, a hetero-bifunctional crosslinker containing N-hydroxysuccinimide (NHS) ester and maleimide groups to approximately 7.35% of available SH groups of the macromer. (**Figure 4A, B**) The modified four-arm NHS-PEG-SH was then mixed in equal volume with guest-host functionalized 4-arm PEG-MAL and emulsified in mineral oil. The resulting NHS-functionalized microgels were then incubated with Armenian hamster anti-mCD3ε (clone 145-2C11) that has previously been shown to reverse T1D in NOD mice. [25, 26] The NHS ester on the microgel surface reacted with primary amines αCD3 to covalently tether the antibody to the microgel surfaces. To demonstrate control over this αCD3 conjugation strategy, we varied the molar amount of Sulfo-SMCC reacted with PEG-SH and detected the amount of αCD3 conjugation by microscopy using a fluorescently-labeled anti-Armenian-hamster secondary antibody. We observed strong uniform conjugation of αCD3 on the surfaces of the microgels (**Figure 4C**). The amount of conjugated αCD3 directly correlated with the molar concentration of Sulfo-SMCC reacted with the PEG-SH, demonstrating the ability to control the dosing and surface availability of αCD3 (**Figure 4D**). Release kinetics of surface-conjugated αCD3 from Sulfo-SMCC microgels at 37°C showed approximately 4% antibody release over three days, demonstrating covalent attachment **(Figure 4E).**

**Figure 4:**
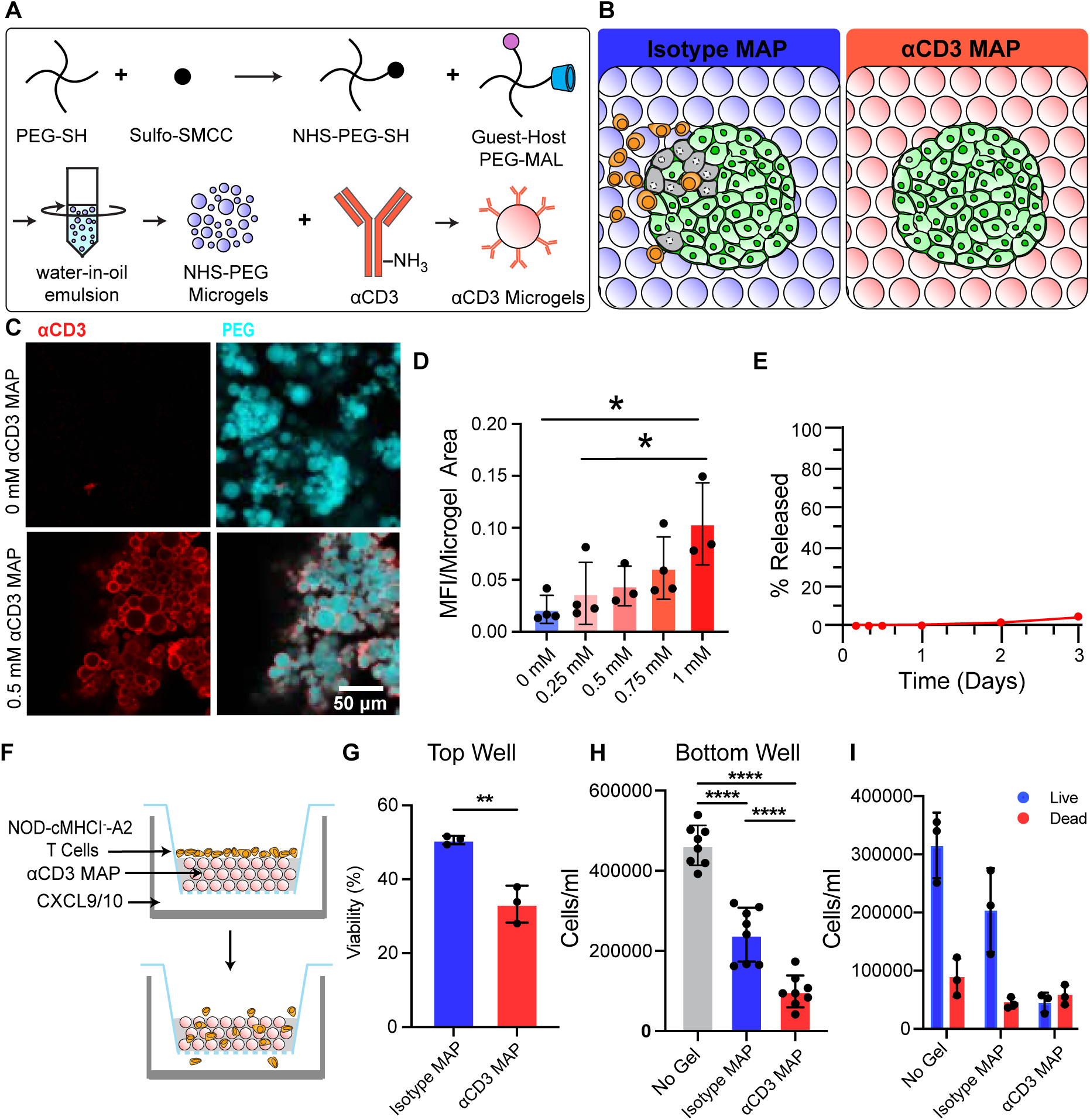
Synthesis of αCD3 microgels to control T cells. (A) Schematic: generation of αCD3 MAP. Four-arm PEG-SH is reacted with a hetero-bifunctional crosslinker, Sulfo-SMCC. The functionalized macromer is then mixed with 4-arm PEG-MAL and made into microgels through a water-in-oil emulsion. Washed microgels are then surface labeled with αCD3 or an isotype control antibody. (B) Schematic: effect of microgels on T cell migration to islet grafts. (Left) Isotype control microgels are permissive to the migration of T cells towards transplanted islets. (Right) αCD3 MAP prevents the migration of T cells towards transplanted islets. (C) Confocal image of guest-host MAP with and without αCD3 (0.5 mM) stained with a fluorescently labeled anti-Armenian hamster secondary antibody. PEG was labeled with Alexa Fluor 647 C5-maleimide during microgel fabrication. (D) Quantified mean fluorescence intensity (MFI) of αCD3 per unit area microgel for each Sulfo-SMCC molar concentration. (n = 3-4) (E) Percent release of αCD3 (0.5 mM) from microgels in PBS over 3 days. (F) Schematic of the experiment, activated T cells from a diabetic NOD-cMHCI⁻-A2 mouse are placed on top of guest-host MAP with or without αCD3 in a transwell insert and mCXCL9 and mCXCL10 are loaded into the bottom well to create a chemotactic gradient. (G) Viability of T cells within microgels in the top well after 18 hours. (H) Total number of T cells migrated to the bottom well after 18 hours. (I) Proportion of live and dead populations in the bottom well after 18 hours. Data analyzed by student’s t-test and one-way ANOVA. N = 7. Mean ± SD.

The functional activity of the conjugated antibody was assessed using a transwell migration assay of isolated, activated CD3^+^ T cells from a diabetic NOD-cMHCI^-^-A2 mouse toward a CXCL9/10 chemotactic gradient through αCD3 MAP (**Figure 4F**). Cells that remained within the αCD3 MAP had lower viability compared to the isotype MAP (**Figure 4G**). Fewer T cells migrated through αCD3 MAP to the bottom well compared with isotype or no-gel controls and had a higher population of dead cells (**Figure 4H, I**). These results indicate that αCD3 MAP can arrest migration of activated T cells, suggesting its potential to block infiltration of autoreactive T cells towards transplanted beta cells *in vivo*.

### αCD3 MAP improves survival of sBCs in a humanized model of type 1 diabetes

To investigate the immunoprotective effects of the αCD3 MAP, we generated sBCs through a direct differentiation process from human Mel1^INS-GFP^ human pluripotent stem cells (**Supplementary Figure 1A**). sBCs at day 23 present as spheroids, express a GFP reporter under control of the insulin promoter (pINS.GFP). Quantification by flow cytometry for expression of the key pancreatic cell marker C-peptide, a byproduct of endogenous insulin synthesis, revealed efficient ∼ 55% sBC generation, consistent with previous reports (**Supplementary Figure 1B, C**). [27–32] Day 23 sBCs were dissociated into single cells, filtered, and cryopreserved. [27] Upon thawing, clusters were reaggregated and cultured in a beta cell maturation media for 5 days.

To model autoimmune rejection of transplanted human sBCs, we employed a humanized-HLA model combining NOD-cMHCI^-^-A2 and NSG-HLA-A2/HHD mice. We confirmed this model by isolating T cells from diabetic NOD-cMHCI^-^-A2 mice, then activating and expanding them before adoptive cell transfer (ACT) into NSG-HLA-A2/HHD recipients transplanted with constitutive firefly-luciferase expressing sBCs. Adoptively transferred T cells effectively infiltrate and destroy both endogenous islets and grafted sBCs, both expressing HLA-A2 on their cell surface, recapitulating critical aspects of autoimmune pathology (**Supplementary Figure 2A-C**). This system provides a reproducible platform to evaluate immunomodulatory biomaterials and immune evasion strategies for beta cell replacement in autoimmune diabetes in a humanized-HLA model.

To investigate the protection of sBCs by αCD3 MAP from a T-cell mediated attack, we transplanted 600 sBC clusters embedded in αCD3 MAP and isotype control MAP into the subcutaneous space of an NSG-HLA-A2/HHD mouse (**Figure 5A**). Following ACT of activated NOD-cMHCI⁻^/^⁻A2 T cells, grafts were monitored via bioluminescence imaging for engraftment and viability over 45 days post ACT (**Figure 5A, B**). Grafts containing αCD3 MAP maintained a detectable signal post ACT, whereas isotype control microgels exhibited loss of signal by day 33. At day 45 post ACT 85% of αCD3 MAP grafts remained detectable compared to 50% of the isotype control, demonstrating a protective effect of αCD3 MAP (**Figure 5C**). Longitudinal blood glucose monitoring showed that transferred T cells trafficked both to the subcutaneous graft and the pancreas, leaving 25% of guest-host MAP mice and 80% of αCD3 MAP mice diabetes-free by day 45 post ACT (**Figure 5D**). Analysis of human C-peptide and mouse C-peptide revealed significantly higher human and mouse C-peptide post-ACT in the αCD3 MAP mice compared to the guest-host MAP control (**Figure 5E, F**). Explanted grafts demonstrated greater numbers of human-nuclear-antigen^+^ (HNA^+^) and insulin^+^ cells remaining in αCD3 MAP compared to the guest-host MAP control (**Figure 5G**).

**Figure 5:**
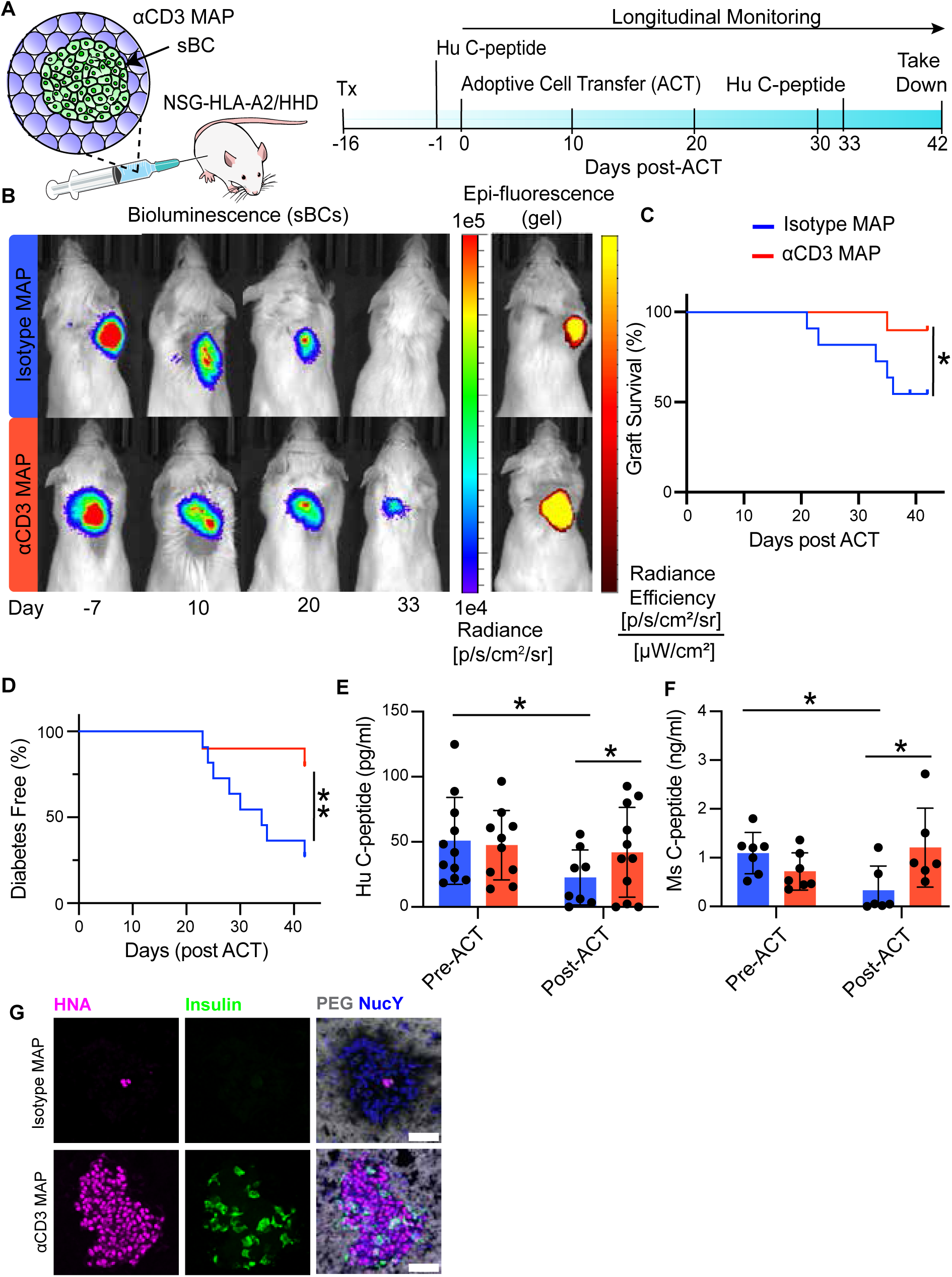
αCD3 MAP improves survival of sBCs in a humanized autoimmune model of type 1 diabetes. (A) Schematic of study. On day-16, 800-1200 sBCs were injected in 200-400 μl of αCD3 MAP or isotype control MAP in the dorsal subcutaneous space of an NSG-HLA-A2/HHD mouse. On day 0, all mice received an adoptive cell transfer of 1e6 activated diabetogenic NOD-cMHCI⁻-A2 T cells. Grafts were then monitored via IVIS over the next 42 days. (B) (Left) Longitudinal monitoring of the sBC graft by transdermal bioluminescence detection via IVIS on day-7, 1, 10, 20, 33, and 42. (Right) Monitoring of fluorescent αCD3 MAP and isotype MAP by fluorescent intensity detection via IVIS on day-7. PEG was labeled with Alexa Fluor 647 C5-maleimide during microgel fabrication. (C) Survival curve of the total flux measured from the sBC grafts (N = 10 mice per group). (D) Percent diabetes free days post-adoptive cell transfer. (E) Human C-peptide and (F) mouse C-peptide measured pre-ACT (Day - 7) and post-ACT (day 33) (G) sBC graft viability stained for HNA (magenta), Insulin (green), PEG (white), and NucY (blue). Scale bar = 50 µm. Data analyzed by nested two-way ANOVA. Survival curves analyzed by Kaplan-Meier Survival Analysis. *p < 0.05, **p < 0.01.

Isotype control guest-host MAP grafts had more infiltrating CD3^+^ T cells surrounding the sBC graft with little to no remaining HNA^+^ or insulin^+^ cells (**Figures 5G, 6A, B**). Flow cytometric analysis of the collected grafts showed no significant differences in the relative frequency between CD4^+^ and CD8^+^ T cells within the graft, indicating the αCD3 MAP was effective on both CD4^+^ and CD8^+^ T cells at the graft site (**Figure 6C**). To understand the effect of αCD3 MAP systemically, we collected the draining lymph nodes and analyzed the T cell compartment by flow cytometry. αCD3 MAP grafts had significantly smaller proportions of circulating CD4^+^ T cells; however, there was no difference in the anergy markers CD73 and FR4 between the two groups. There were also no differences between proportions of CD8^+^ T cells or their anergic markers (**Figure 6I)**. In the pancreas, there were significantly more islets per mouse and brighter insulin staining in αCD3 MAP (**Figure 6J**). There were significantly fewer infiltrating CD3^+^ T cells in the pancreas for αCD3 MAP treated mice (**Figure 6K, L**). Together, these results support αCD3 MAP deletion or arrest of draining diabetogenicT cells, thereby limiting autoimmune destruction and preserving sBC viability.

**Figure 6:**
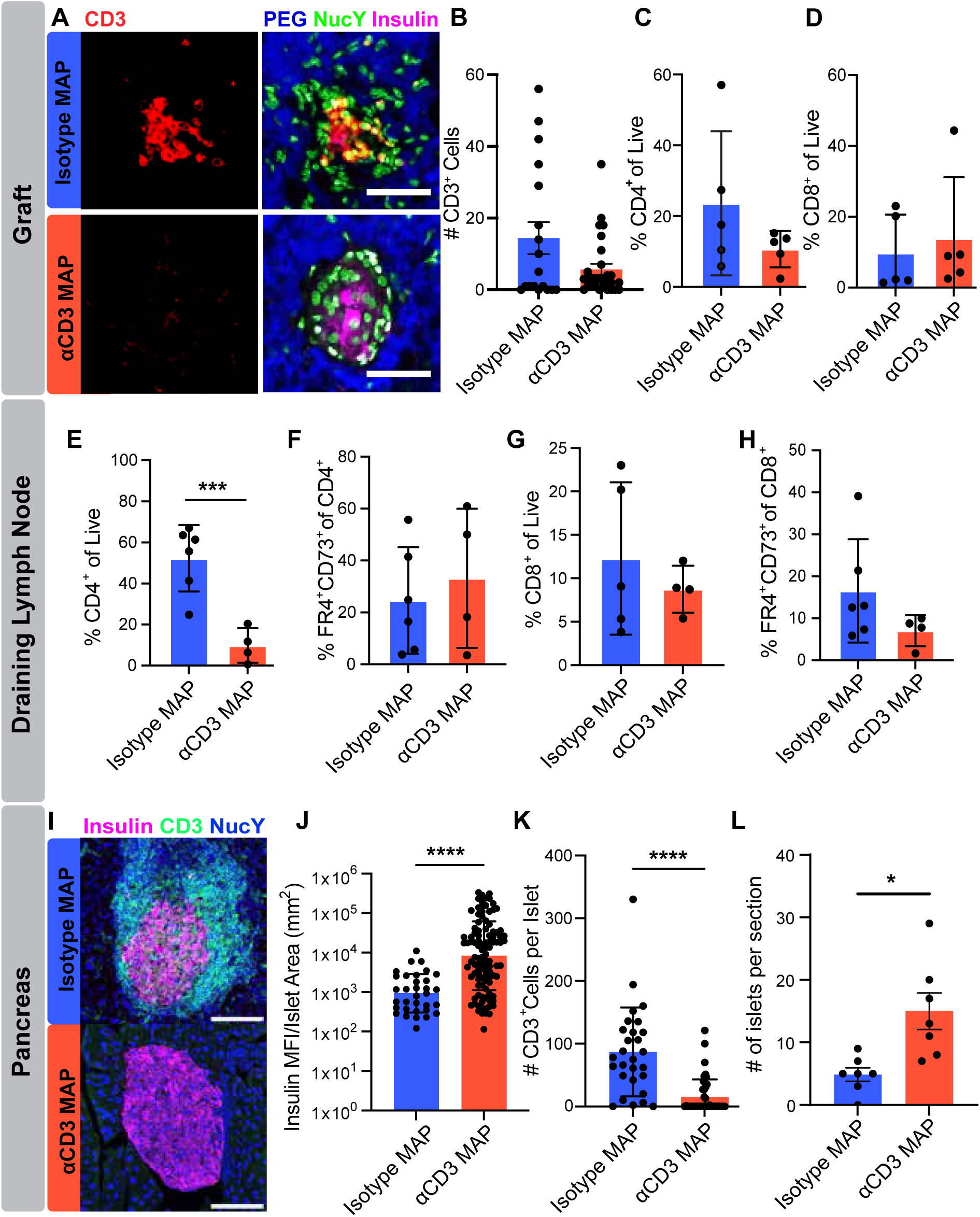
αCD3 MAP reduces T cell infiltration of grafted sBCs and native islets in the pancreas. (A) Confocal image of day 42 explanted graft immunostained for insulin (beta cells) and CD3 (T cells). Nuclear yellow was used as a nuclear counterstain. PEG was labeled with Alexa Fluor 647 C5-maleimide during microgel fabrication. Scale bar = 50 μm. (B) Quantification of CD3^+^ T cells surrounding sBC grafts in tissue sections. (C) Percentage of CD4^+^ and (D) CD8^+^ T cells within microgel graft area analyzed by flow cytometry. (E-H) Flow cytometry analysis of draining lymph nodes of αCD3 MAP or isotype control MAP treated mice. (E) Percentage of CD4^+^ of live (F), percentage of FR4^+^CD73^+^ of CD4^+^, (G) Percentage of CD8^+^ of live, and (H) Percentage of FR4^+^CD73^+^ of CD8^+^. (I) Representative confocal images of islets in the pancreas of αCD3 MAP and isotype MAP treated mice, immunostained for insulin and CD3, and counterstained with Nuclear Yellow. Scale bar = 100 μm (J) Quantification of insulin MFI per unit insulin^+^ area in pancreas. (K) Quantification of CD3^+^ T cells surrounding islets in the pancreas. (L) Number of islets quantified per section. (N = 3 sections per mouse). Data analyzed by student’s t-test. N = 4-5 mice for flow cytometry. N = 10 mice for pancreas quantification. *p < 0.05, ** p < 0.01, ***p < 0.001, ****p < 0.0001.

## Discussion

This study aimed to generate a localized immunomodulatory niche using αCD3 MAP for enhancing the engraftment and survival of transplanted sBCs in the context of autoimmune diabetes. Our findings demonstrate that guest-host MAP hydrogels provide a mechanically resilient and cytocompatible scaffold that supports vascular integration in the subcutaneous space and serves as a framework for the tethered presentation of immunomodulatory molecules such as αCD3 to attenuate autoreactive T cell responses.

Compared to PEG microgels, guest-host MAP exhibited enhanced mechanical resilience and recovery following high-strain deformation, supporting their use as injectable materials capable of withstanding the dynamic stresses of transplantation. Importantly, guest-host MAP hydrogels did not elicit cytotoxic effects on murine islets *in vitro*, suggesting that the guest-host interactions themselves do not compromise cell viability. *In vivo*, guest-host interactions were advantageous for minimizing the foreign body response to the implant and increasing vascularization. While both guest-host MAP and PEG microgels exhibited minimal fibrotic encapsulation, guest-host MAP better supported vascularization. The guest-host MAP hydrogel likely supported higher vascularization than the PEG microgels because of the combination of interconnected, cell-scale porosity with moderately increased stiffness and network stability that has been shown to favor angiogenic sprouting and vessel stabilization while resisting fibrotic encapsulation. [24, 33–35] These findings highlight the importance of microporosity and dynamic guest-host interactions in creating an immune-permissive environment that supports neovascularization.

Robust vascular integration is particularly critical in the subcutaneous niche, which is otherwise poorly vascularized and has historically been inhospitable to beta cell grafts. Recent advances have demonstrated success in islet transplantation to the subcutaneous space using strategies such as amino-acid-rich collagen matrix, macroencapsulation devices, and engineered scaffolds. [36–42] The advantages of our approach are its simplicity and injectability, ability to be delivered in a single step with no post-injection annealing needed, and it does not require loading of the scaffold with supplemental nutrients or recombinant growth factors to achieve islet engraftment in the subcutaneous space.

Functionally, guest-host MAP hydrogels supported the survival and glucose-regulating capacity of transplanted syngeneic islets in a streptozotocin-induced diabetes model. Primary islets transplanted subcutaneously and embedded within guest-host MAP restored normoglycemia by eight weeks and demonstrated glycemic control kinetics comparable to non-diabetic controls. Histological analysis confirmed that engrafted islets became vascularized and integrated into the host tissue, consistent with the improved metabolic outcomes observed *in vivo*. By contrast, islet-only controls failed to establish stable grafts, underscoring the necessity of a supportive biomaterial scaffold for long-term function in the subcutaneous space.

By conjugating αCD3 to the hydrogel surface, we created a localized, sustained immunomodulatory interface that engages and functionally impairs autoreactive T cells. *In vitro* assays confirmed that αCD3 MAP reduces T cell viability and prevents direct T cell migration toward chemokine gradients, demonstrating that the tethered antibody retains its CD3-binding capacity when covalently tethered to the microgel matrix. Importantly, release kinetics studies indicated minimal antibody release, supporting the notion that immune modulation occurs locally at the graft site rather than systemically.

*In vivo*, we leveraged an HLA-humanized adoptive cell transfer model of autoimmune diabetes to evaluate the immunoprotective function of αCD3 MAP in a controlled manner. There are limited options for testing T1D-like autoimmune rejection of human beta cells *in vivo*. The use of humanized mice reconstituted with immune systems from T1D donors has begun to address this issue. However, there are major challenges with these models: mice hosting human immune systems are expensive and difficult to generate; human immune cells have difficulty with survival and trafficking to the peripheral tissues in mice; the host animals tend to develop graft-versus-host disease within 30 days; and there are concerns with distinguishing allo-immunity from T1D-like autoimmunity. [43] Here, we developed a novel approach to modeling T1D autoimmunity *in vivo*, where mouse diabetogenic CD8^+^ T cells natively recognize human HLA by knocking out the mouse MHC class I and expressing human HLA class I. Thus, when adoptively transferred into a naïve NSG host harboring the same human HLA transgene, the mouse T cells traffic normally, transfer diabetes, recognize the human beta cell graft, and do not create graft versus host disease. A limitation of our model is the need for a shared T1D antigen between human and mouse beta cells. Islet-specific glucose-6-phosphatase catalytic subunit-related protein (IGRP), IGRP_265-273_, is one such major T1D CD8^+^ epitope that drives disease in both humans and NOD mice and is likely important in our model. [44]

Another limitation of our model is that the human beta cells express other xeno-antigens that could trigger an adaptive immune response. We delayed the xeno-recognition by using naïve NSG mice as the host animal, where we demonstrated lack of rejection of the human sBC graft until after the adoptive transfer of activated diabetogenic T cells. However, we could not show indefinite sBC survival in this model because of the eventual development of an adaptive immune response, likely against xeno-antigens. Nevertheless, our model is among the first to demonstrate T1D-like autoimmunity directed against human beta cells in an HLA-matched manner in the *in vivo* setting, combined with a biomaterials strategy to mitigate T1D autoimmunity against replacement beta cells.

This approach allowed us to precisely target diabetogenic T cells while avoiding confounding contributions from B cells, NK cells, or other innate immune compartments. Under this stringent challenge, sBCs transplanted with αCD3 MAP exhibited improved survival compared to isotype controls, with preserved graft signal, increased HNA^+^ and insulin^+^ cell retention, and higher human C-peptide levels. Moreover, αCD3 MAP protected not only the local graft but also preserved endogenous pancreatic islets, suggesting a systemic benefit through deletion or arrest of autoreactive T cells that were attracted to the graft site. Flow cytometry analysis supported this interpretation, revealing reduced proportions of CD4^+^ T cells in draining lymph nodes without clear induction of an anergic phenotype, consistent with αCD3-mediated depletion or functional inactivation.

These findings highlight several important mechanistic insights. First, localized presentation of αCD3 within a supportive biomaterial niche can limit autoreactive T cell infiltration and function without broad systemic immunosuppression. Second, the protection of both transplanted sBCs and endogenous pancreatic islets indicates that αCD3 MAP may act as a sink for autoreactive T cells, promoting their deletion during trafficking through the graft site. Third, the dual role of MAP, serving as both a physical scaffold for vascularization and a biochemical platform for tethered immunomodulation, addresses two critical and often competing challenges in cell replacement therapy of graft survival in poorly vascularized transplant sites versus the resistance to autoimmune destruction.

While these results are promising, several limitations warrant consideration. Although robust vascularization is seen in the graft site after several weeks, we hypothesize that the delay in re-vascularization contributes to some degree to the beta cell loss in the time period immediately after injection. In addition, the αCD3 antibody used throughout these studies still contains the Fc region, as opposed to the Fc non-binding variations that are now being employed in clinical trials. [17] Previously, the presence of this Fc region has been shown to promote T cell proliferation and cytokine production through the engagement of the Fc region with Fc receptors (FcR) on monocytes and macrophages. [45, 46] However, this effect was shown only for the soluble form of the antibody, and covalently immobilized antibodies may be less able to bind FcR. Here, we used a mouse antibody to interface with mouse immune cells. To increase translatability in future studies, the humanized mAb Teplizumab will be assessed in fully humanized models to evaluate the durability of protection and potential for long-term immune escape.

In summary, we demonstrate that αCD3 MAP arrests autoreactive T cell infiltration, preserves graft viability, and protects endogenous islets from immune-mediated destruction in a stringent subcutaneous HLA-humanized adoptive cell transfer model. This work introduces a novel strategy for localized, durable immune modulation in type 1 diabetes, offering an alternative to systemic immunosuppression and advancing the clinical feasibility of beta cell replacement therapies.

## Methods

### Guest-host MAP and PEG Microgel Fabrication

Guest-host MAP microgels were synthesized by dissolving four-arm PEG-MAL (Laysan Bio #4-ARM-PEG-MAL-20k, 20kDa) in 1X phosphate buffered saline (PBS) with 1% HEPES at pH 5.4 (31.4 mg/ml, 1.5 mM). The reduced pH helps to slow down the crosslinking reaction, which at neutral pH is so quick it hinders mixing and handling. [47] At pH 5.4, the gelation of PEG-4MAL with PEG-4SH takes approximately 34 seconds. [48] Adamantane-thiol (Sigma Aldrich #659452) was dissolved in 1X PBS with 1% HEPES and 10% DMSO at pH 5.4 (0.084 mg/ml, 0.05 mM), and mono-(6-mercapto-6-deoxy)-β-cyclodextrin (Zhiyuan Biotechnology) was dissolved in 1X PBS with 1% HEPES at pH 5.4 (0.56 mg/ml, 0.05 mM). Adamantane and β-cyclodextrin were added dropwise to the PEG-MAL macromer at a 1 adamantane:1 β-cyclodextrin molar ratio and reacted for 30 minutes at 25°C. Four-arm PEG-SH (Laysan Bio #4-ARM-PEG-SH-20K, 20kDa) was dissolved in 1X PBS with 1% HEPES at pH 5.4 (28.5 mg/ml, 1.4 mM) and reacted with a trace amount of Alexa Fluor 647-maleimide for 5 minutes at 25°C to aid in microgel visualization. PEG-SH was added to the functionalized PEG-MAL macromer at a 1:1 volume ratio, quickly pipetted up and down several times to mix thoroughly, then transferred to a 30x volume of mineral oil with 2% vol/vol span 80 surfactant in a 50 ml conical tube. The tube was immediately vortexed for 30 seconds to generate an emulsion, then allowed to finish crosslinking for 1 hour at 25°C on a rocker plate to generate microgels with a final weight percentage of ∼ 6% wt/vol. Unfunctionalized PEG microgels were synthesized by dissolving PEG-MAL macromer in 1X PBS with 1% HEPES at pH 5.4 (30.4 mg/ml, 1.4 mM). PEG-SH was dissolved in 1X PBS with 1% HEPES at pH 5.4 (29.6 mg/ml, 1.4 mM) and reacted with a trace amount of Alexa Fluor 647-maleimide for 5 minutes. PEG-SH was added dropwise to the PEG-MAL macromer at a 1:1 volume ratio, and microgels were formed by emulsion the same as above. For all groups, microgels were centrifuged at 3,000 x *g* for 5 minutes and washed three times with 0.3 % Triton X-100 in 1X PBS at pH 7.2 and twice with 1X PBS.

### Microgel Size Characterization

Microgel diameter was measured by imaging several batches of microgels using confocal microscopy. The images were processed using a gaussian filter (radius = 2) and automatically segmented into regions of interest to determine the microgel diameter using StarDist in the FIJI distribution of ImageJ [49, 50]. The Feret’s diameter of each microgel was considered as the diameter. The particle size data were then analyzed by frequency distribution with a bin size of 5 µm, the average diameter and standard deviation were used to determine the PDI (**Equation 1**), where σ is the standard deviation and <D> is the average diameter. [51]

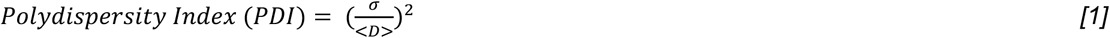

### Rheological Characterization

Rheological measurements were performed on an Anton Paar MCR 702 rheometer with a 20mm sandblasted plate on plate configuration with 0.5 mm gap height at 25°C. To load samples on the rheometer, approximately 1 mL of microgels were placed on the bottom plate. The top plate was lowered to the correct gap height, and microgels that did not fit into the geometry were removed with a spatula. Oscillatory frequency sweeps were performed from 0.01 Hz to 10 Hz at 1% strain. Oscillatory strain cycle and recovery experiments were conducted by alternating 1% for 120s and 500% strain for 60s over three periods at 1 Hz. Unidirectional shear rate sweeps were performed between 100 1/s and 0.01 1/s shear rate (*γ),* and measuring shear stress (*σ*). A linear regression was performed to determine the zero-frequency limit and resultant yield stress of the microgels on GraphPad.

### C57BL/6J islet isolation

Naïve C57BL/6J mice aged at least 16 weeks were deeply anesthetized with isoflurane and euthanized by cervical dislocation. A midline laparotomy was performed, and the liver was lifted over the diaphragm to visualize the underlying organs. The pancreas was perfused with 0.6 mg/mL collagenase by ligating the hepatopancreatic ampulla with a nylon suture and injecting the digestion solution through the common bile duct. After inflation of the pancreas, the whole organ was carefully removed and placed on ice. Pancreata were incubated in a 37°C water bath for 17 minutes before stopping digestion with ice cold stop solution (88% HBSS, 9% FBS, 2% HEPES, 1% penicillin-streptomycin) and washing with serum-free solution (97% HBSS, 2% HEPES, 1% penicillin-streptomycin). For co-culture studies, islets were handpicked; for transplants, islets were separated from exocrine tissue using a Histopaque-1119 (Sigma Aldrich #11191) gradient. Before further use, islets were cultured for 24-48 hours in a 37°C, 5% CO_2_ incubator in mouse islet medium (88% RPMI 1640, 10% FBS, 1% Glutamax, and 1% penicillin-streptomycin).

### Islet Viability in Microgels

20 Isolated C57BL/6 islets were mixed with 100 μl of gel, 200 μl of mouse islet media was placed on top of the gel and islet mixture and incubated at 37°C for seven days. Media was switched every two days, samples were stained with CalceinAM (5 μM), Ethidium homodimer-1 (EthD-1) (2 μM), and Hoescht 34580 (10 μg/mL) (Invitrogen #H21486) for 30 min at 37°C and imaged on days one, five, and seven to assess viability. 4-5 islets per group were imaged on a confocal microscope with a 20x air objective. Numbers of cells were quantified by the StarDist package in FIJI, and viability was determined by subtracting the proportion of dead cells from the total. [49, 50]

### Subcutaneous microgel implantation

C57BL/6J mice were anesthetized with 1.5-2% isoflurane, shaved, and cleaned with alternate washes of sterile saline and 2% chlorhexidine. Subcutaneous injection of meloxicam was administered as a post-operative analgesic. 20 μl of guest-host MAP and PEG microgels were loaded into the back of a sterile 1 mL syringe, and islets were loaded into the center of the microgels using a pipette. The back of the syringe was replaced and used to compact the gel and islet mixture to the bottom of the syringe and fitted with a 20G ½” needle. Guest-host and unfunctionalized microgels with islets were injected into the upper left and lower left quadrant of the dorsum, respectively. Grafts were left to engraft for 31 days before being removed and fixed in 3.2% paraformaldehyde at 4°C overnight and processed for histology as described below.

### Streptozotocin model of hyperglycemia

C57BL/6J mice aged 13-16 weeks received a single 160 mg/kg dose of streptozotocin (STZ, Sigma Aldrich #S0130) intraperitoneally to induce diabetes. STZ was injected immediately after dissolution in ice cold 0.9% normal saline to maximize potency. Following STZ administration, all mice were monitored daily for the onset of hyperglycemia and given access to 10% wt/vol sucrose water *ad libitum* for the first three days to prevent hypoglycemia. Mice with at least two consecutive blood glucose readings above 350 mg/dL (collected via tail vein puncture) were considered candidates for transplantation.

### Subcutaneous islet transplantation

Guest-host MAP was swelled in 10x vol/vol serum-free medium (RPMI 1640, 10 mM Glutamax, 10 mM cysteine) overnight at 4°C on a rocker or for two hours at 37°C prior to transplantation. On the day of transplantation, islets were hand-counted into aliquots of 400 islets, washed once with PBS, and resuspended in serum-free medium (same formulation). The formulation was chosen to minimize xenogeneic immune responses and mitigate islet ischemia. Diabetic mice were anesthetized with 1.5-2.5% isoflurane, and the lower left quadrant of the dorsum was shaved. Mice received a 15 mg/kg dose of meloxicam subcutaneously in the upper right quadrant of the dorsum, to avoid the transplant site, with two additional doses given at 24-hour intervals after the procedure. The site was then cleaned three times with alternating washes of 2% chlorhexidine and 0.9% normal saline. To transplant the islets, 200 µL of guest-host MAP was first aseptically loaded into the back of a 1 mL syringe. Then, the entire contents of one islet aliquot were aspirated in a 200 µL pipette tip, allowing time for the islets to settle before dispersing into the center of the gel. The syringe plunger was then replaced and used to compact the gel/islet mixture. After fitting the syringe with a 26G ½” needle, the entire contents of the syringe were injected in the subcutaneous space.

### Longitudinal monitoring of syngeneic islet grafts

Blood glucose was monitored daily via tail vein puncture until day 7 post-transplantation, or until amelioration of severe diabetes (blood glucose >400 mg/dL). During this time, mice also received daily subcutaneous injections of 0.5-1 units of Lantus long-acting insulin (Sanofi). Blood glucose was monitored three times per week thereafter, with no exogenous insulin delivery. At days 28, 42, 56, and 70 post-transplantation, mice were fasted for six hours (>24 hours after the last exogenous insulin dose) before challenge with 1 g/kg glucose solution. Blood glucose values for the intraperitoneal glucose tolerance test (IPGTT) were collected at 0, 5, 10, 20, 30, 60, and 90 minutes after injection. Grafts were removed for histology at day 84 post-transplantation, and animals were euthanized.

### aCD3 MAP Fabrication

Four-arm PEG-MAL (20kDa) and four-arm PEG-SH (20kDa) were dissolved in 1X PBS with 1% HEPES at pH 5.4 (34.5 mg/ml, 1.7 mM). Sulfo-SMCC was dissolved in 1X PBS at pH 5.4 (0.21 mg/ml, 0.5 mM) and reacted with PEG-SH for 30 minutes at 25°C. Adamantane-thiol and mono-6-mercapto-β-cyclodextrin were functionalized to PEG-MAL as described above. Sulfo-SMCC functionalized PEG-SH was added to the functionalized PEG-MAL macromer at a 1:1 volume ratio; microgel generation and washing steps were completed as described above. Microgels were suspended in a 1:2 volume ratio with 1X PBS, and 200 μl of 0.2 mg/ml anti-mouse CD3ε (Biolegend, 145-2C11) or Armenian Hamster IgG isotype control and allowed to react for 16 hours. Post-incubation, microgels were centrifuged at 3,000 x *g,* and the supernatant was removed and washed with 1X PBS.

### Human stem cell culture and sBC differentiation

Undifferentiated human pluripotent stem (hPSC) Mel1^INS-GFP^ reporter cells [52] engineered to express a firefly luciferase construct were maintained on hESC qualified Cultrex (Biotechne #3434-005-002) in mTeSR+ media (STEMCELL Technologies #05826). Differentiation to stem cell-derived beta-like cells (sBC) was carried out in suspension-based, magnetic stirring platforms (Reprocell #ABBWVS03A-6, #ABBWVDW-1013, #ABBWBP03N0S-6) as described [53, 54]. Briefly, 90% confluent hPSC cultures were dissociated into a single-cell suspension by incubation with TrypLE (Gibco #12-604-021). Dissociation was halted with mTeSR+ media, and cells were counted using a Countess 3 cell counter (ThermoFisher Scientific), followed by seeding 0.5 × 10^6^ cells/ml in mTeSR+ media supplemented with 10 μM ROCK inhibitor in bioreactors. 3D sphere formation was performed for 48-72 hours. Differentiation media was changed daily by letting spheres settle by gravity for 3-5 min. Most supernatant was removed by aspiration; fresh media was added, and bioreactors were placed back on the stirrer system. sBC differentiation and cryopreservation were based on our published protocol [27, 55] with modifications as outlined below. Differentiation medias are as follows: induction of definitive endoderm differentiation using **d1 media** [RPMI containing 0.2% FBS, 1:5,000 ITS (Gibco #41400-045), 200 ng/ml Activin A (R&D Systems #338-AC-01M), and 3 μM CHIR99021 (STEMCELL Technologies #72054)] and **day 2-3**: RPMI containing 0.2% FBS, 1:2,000 ITS, and 100 ng/ml Activin A; **d4-5**: RPMI containing 2% FBS, 1:1,000 ITS, and 50 ng/ml KGF (Prepotech #100-19-1MG); **d6-7**: DMEM with 4.5 g/l D-glucose (Gibco #11960-044) containing 1:50 N-21 MAX (Biotechne #AR008), 1:100 NEAA (Gibco #11140-050), 1mM Sodium Pyruvate (Gibco #11360-070), 1:100 GlutaMAX (Gibco #35050-061), 3 nM TTNPB, (R&D Systems #0761), 250 nM Sant-1 (R&D Systems #1974), 250 nM LDN (STEMCELL Technologies #72149), 30 nM PMA (Sigma Aldrich #P1585-1MG), 50 μg/ml 2-phospho-L-ascorbic acid trisodium salt (VitC) (Sigma #49752-10G); **d8-9**: DMEM containing 1:50 N-21 MAX, 1:100 NEAA, 1 mM Sodium Pyruvate, 1:100 GlutaMAX, 100 ng/ml EGF (R&D Systems #236-EG-01M), 50 ng/ml KGF, and 50 μg/ml VitC; **d10-15**: DMEM containing 1:50 N-21 MAX, 1:100 NEAA, 1 mM Sodium Pyruvate, 1:100 GlutaMAX, 10 μg/ml Heparin (Sigma #H3149-250KU), 2 mM N-Acetyl-L-cysteine (Cysteine) (Sigma #A9165-25G), 10 μM Zinc sulfate heptahydrate (Zinc) (Sigma #Z0251-100g), 1x BME, 10 μM Alk5i II RepSox (R&D Systems #3742/50), 1 μM 3,3’,5-Triiodo-L-thyronine sodium salt (T3) (Sigma #T6397), 0.5 μM LDN, 1uM Gamma Secretase Inhibitor XX (XXi) (AsisChem #ASIS-0149) and 1:250 1 M NaOH to adjust pH to ∼7.4. For experiments presented in Figures 4, 5, 6, 7, 8 and Supplementary Figures 4, 5, 7 and 8 10μM ROCKi (RnD systems #1254/50) was added; **d16-23**: CMRL (Gibco #11530-037) containing 1:50 N-21 MAX, 1:100 NEAA, 1:100 GlutaMAX, 10ug/ml Heparin, 2mM Cysteine, 10 μM Zinc, 1x BME, 1 μM T3, 50ug/ml VitC, 1:1000 Trace Elements A (Corning # 25-021-CI), 1:1000 Trace Elements B (Corning # 25-022-CI) with or without 10 μM Alk5i II RepSox and 1:250 NaOH to adjust pH to ∼7.4. All medias with exception of mTeSR+ also contained 1x PenStrep.

### Cryopreservation and thawing of hPSC-derived beta-like cells

Day 23 sBC were dissociated into single cells and cryopreserved as described. [27, 53] Briefly, cells were quenched with 2% FBS in PBS and filtered using a cell strainer into FACS 5 ml tubes. Cells were counted using a Countess 3 cell counter (ThermoFisher Scientific) and resuspended at 3 x 10^6^ cells/100 μl of CryoStor® CS10 (StemCell Technologies). Cells were cryopreserved overnight before transfer to liquid nitrogen for long-term storage. For thawing, 1 ml of warm sBC media (d16-23 media described above) was added to the thawed cryovial dropwise before the entire volume was transferred into 5 ml of sBC media, counted, and seeded in Aggrewell 800 plates to generate clusters with 3,000 cells/cluster. After 24 hours, a partial media change was done to remove any debris. Fully formed clusters were generated after 48-72 hours.

### Autoimmune diabetes model and Adoptive Cell Transfer

The mouse strain NOD-cMHCI^-^-A2 (Jax Stock No: 031856) was provided by Dr. Dave Serreze at The Jackson Laboratory, and commercially available NSG-HLA-A2/HHD mice (Jax Stock No: 014570) were ordered from The Jackson Laboratory. Mice were maintained in a vivarium managed by the University of Florida Institutional Animal Care Services, and genotype was confirmed by PCR using Transnetyx commercial genotyping services. Splenocytes from NOD-cMHCI^-^-A2 were collected and frozen after blood glucose readings over 250 mg/dl. Splenocytes were thawed and isolated for CD3^+^ T Cells using the EasySep^TM^ Mouse T Cell Isolation Kit (Stemcell Technologies, no.19851). 6-well plates were coated with anti-CD28 and anti-CD3 overnight in 1X PBS at 4°C. Plates were then blocked for 4 hours with splenocyte media at 25°C. Isolated T cells were plated and left to activate for 3 days with 100U/mL of hIL-2. After 3 days, T cells were removed from coated plates and allowed to expand in a T-75 flask for 3 days. Five million activated T cells were injected into the tail vein of the mice for the initial confirmation of the model (Supplementary Figure 1). For assessment of graft viability, one million activated T cells were injected via the tail vein into the mice.

### Transwell migration assay of NOD-cMHCI--A2 towards a chemokine gradient

NOD-cMHCI^-^-A2 T cells were activated and expanded as described above. 200 μL of aCD3 MAP and isotype MAP were packed into the back of a 1mL syringe and injected on top of a transwell insert (8 μm pore size, 24-well plate, Corning #3464). MAP hydrogels were spun down in the transwell insert at 3000 x *g* for 5 minutes to pack the MAP hydrogels in the transwell. 1 x 10^6^ activated and expanded NOD-cMHCI^-^-A2 T cells were placed on top of the gel in the transwell in 400 μL of cRPMI. Additional cRPMI with 0.3 μg/mL of both CXCL9 (mCXCL9/MIG, R&D systems #492-MM) and CXCL10 (mCXCL10-10MP-10/CRG-2, R&D Systems #466-CR) were added into the bottom well. After 24 hours, the number of cells in the bottom well was assessed using 3 measurements on a Countess, assessing viability with a 1:1 dilution in Trypan Blue. The top well was labeled with CalceinAM (5 μM), Ethidium homodimer-1 (EthD-1) (2 μM), and Hoescht 34580 (10 μg/mL) (Invitrogen #H21486) for 30 min at 37°C. Three, 200 μm Z-stacks (Z-step = 2 μm) were taken for each well. The number of live, dead, and total cells was quantified by the StarDist package in FIJI, viability was determined by subtracting the proportion of dead cells from the total. [49, 50]

### Longitudinal Monitoring of the sBC grafts by transdermal bioluminescence

Following transplantation, the sBC graft viability was longitudinally monitored by transdermal bioluminescence. D-Luciferin (Revvity #122799) was prepared at a concentration of 30 mg/ml and filtered through a 0.22 μm Spin-X® centrifuge tube filter (Costar no. 8160) at each timepoint. Mice were anesthetized with 1.5-2% isoflurane, and sterile ocular lubricant was applied. Hair on the right dorsal side of the mice was removed with depilatory cream.

D-Luciferin was injected subcutaneously at 75 mg/kg 15 minutes prior to imaging. Transdermal bioluminescence was captured with an IVIS® Spectrum *In Vivo* Imaging System. For bioluminescence imaging, the system was set to take images without any emission filter (open), and sequential imaging was performed until the signal peaked. Transplants were longitudinally monitored every five days after transplant and every three days after adoptive cell transfer. *In vivo* bioluminescence imaging data was analyzed using PerkinElmer, Inc. Living Image® 4.5.5 software. The total flux (o/s) was estimated with a circular region of interest (ROI) over the graft. A corrected total flux (CTF) was determined by normalizing to the background CTF of the left foot **(Equation 2**).

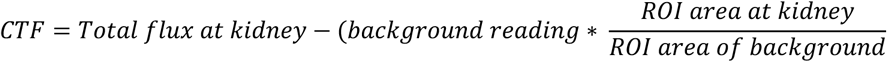

### Serum C-peptide Analysis

For serum C-peptide, mice were fasted for 6 hours, followed by an intraperitoneal injection of 2g/kg body weight glucose in a 25% glucose solution. Peripheral blood was collected via tail puncture at 30 minutes post-injection into anti-coagulant serum tubes. Blood tubes were spun down at 10,000 x g for 10 minutes, and serum was isolated into new tubes. Human C-peptide was quantified using a human specific C-peptide ELISA (ALPCO: 80-CPTHU-CH05). Mouse C-peptide was quantified using a mouse specific C-peptide ELISA (Crystal Chem: 90050).

### Flow Cytometry

Grafts and draining brachial lymph nodes were collected from the animal and placed on cold 1X PBS. Single cell suspensions were made by washing clusters with PBS and incubating with TrypLE or Collagenase-D at 37°C for 12-15 min. Cells were quenched with 2% FBS in PBS and filtered using a cell strainer into FACS 5 ml tubes. Samples were incubated for 10 min on ice with Fc block and subsequently stained with surface markers/live dead staining for 30 min. Samples were fixed with 10% formalin for 10 min at room temperature, then stained in CAS buffer with 0.4% Triton X (CAS-T) overnight at 4°C for intracellular markers. After incubation, the cells were washed and resuspended in FACS buffer for analyses on a 5-laser Cytek Aurora. Analysis was done using FlowJo v10.9 (BD Life Sciences). The following antibodies were used with their respective dilution: PE/Cyanine7 anti-mouse FR4 (Biolegend #125012, 1:100), PerCP/Cyanine5.5 anti-mouse CD73 (Biolegend #127214, 1:100), Brilliant violet 750 anti-mouse CD4 (Biolegend #100467, 1:200), Brilliant violet 510 anti-mouse CD8a (Biolegend #100751, 1:200), PE mouse anti-human Chromagranin A (BD Pharmingen #564563, 1:50), C-peptide mouse monoclonal antibody (OriGene Technologies #BM270, 1:500, self-conjugated using Antibody labeling kit Alexa fluor 488 (Invitrogen #A88062).

### Histology and Immunofluorescence

Dorsal sections containing the gel grafts were fixed with 3.2% paraformaldehyde or 10% formalin at 4°C in 1X PBS overnight, followed by three washes with 1X PBS. The tissues were immersed in 15 % w/v sucrose in PBS overnight and were followed by 30 % w/v sucrose overnight, then snap frozen in Optimal Cutting Temperature (OCT) compound using 2-methylbutane (>99% purity) chilled with liquid nitrogen. Frozen tissue blocks were cut with a cryostat into serial 20 μm sections and mounted on superfrost plus microscope slides. Hematoxylin and eosin stains were performed by the University of Florida Molecular Pathology core and imaged using an Olympus VS200. Using Cell Profiler, images were separated into color images, and the nuclei were identified as objects from the hematoxylin stain. Using the relate objects and find object neighbors module in Cell Profiler, the nearest distance of the nuclei to the surface of the graft was measured. Five sections of each graft were analyzed per mouse.

For immunofluorescence imaging, cryosections were washed three times with 0.3% TX-100 for 5 minutes each. Samples were then blocked and permeabilized in PBS with 0.3 % Triton X-100 + 10% donkey serum for 1 hour at 25°C. Primary antibodies were incubated overnight in PBS with 0.3% Triton X-100 + 1% donkey serum at 4°C. Secondary antibodies were incubated at 1:200 dilutions in PBS with 0.3% Triton X-100 for 1 hour at room temperature. Some sections were stained with Nuclear Yellow (Hoescht S769121) at 1:50 in 1X PBS with 0.3% TritonX-100, then washed with 0.3% Triton X-100 three times before coverslips were mounted with ProLong Gold. The following primary antibodies were used for immunofluorescence staining with their respective dilution: rabbit anti-laminin (Abcam #ab11575, 1:300), goat anti-Col-IV (Abcam #ab6586, 1:40), goat anti-CD31 (Biotechne #AF3628, 1:600), guinea pig anti-insulin (DAKO, #A0452, 1:1000), rabbit anti-HNA (Thermo Fisher Scientific #RBM5-346-P1, 1:200), rat anti-CD3 (Biolegend #100202, 1:50), and rabbit anti-CD8 (Bioss #bs-0648R-TR, 1:200).

### Statistical analysis

Means among three or more groups were compared by a one-way or two-way analysis of variance (ANOVA) in GraphPad Prism 10 software. If deemed significant, Tukey’s post-hoc pairwise comparisons were performed. Means between two groups were compared by a two-tailed Student’s t-test. Experiments with hierarchical data (multiple slides/images from each of multiple independent biological units/mice) where measured by nested t-test. A confidence level of 95% was considered significant. Survival curves were analyzed using a Kaplan-Meier curve and analysis. The statistical test used, error bars, and definition of n are indicated in the individual figure legends.

## Microscopy

MAP hydrogels and tissue sections were imaged on a Leica SP8 confocal laser-scanning microscope using 10×/0.3 and 20×/0.8 numerical aperture Plan-Apochromat air objectives at 1024 × 1024-pixel resolution.

## Author Contributions

AEW, HAR, and EAP conceived and designed the study. AEW, CTM, JMB, HAR, and EAP analyzed and interpreted the results. AEW and EAP wrote the manuscript. AEW, JMB, and AML generated the sBCs. AEW, CTM, and CPS generated and characterized the biomaterial platforms. AEW, CTM, CPS, and JMB conducted animal experiments. AEW, CTM, CPS, and ADG performed histological analysis. All authors discussed the results and commented on the manuscript. All authors contributed to the article and approved the submitted version.

## Acknowledgements

The authors would like to acknowledge Dr. Dave Serreze at The Jackson Lab for providing the associated mouse models. The authors thank Anton Paar for the use of the Anton Paar 702 rheometer through their VIP academic research program and Dr. Thomas Angelini for the use of his lab space.

