## Supplementary Figure for "Anti-CD3 microporous annealed particle hydrogel protects stem cell derived beta cells from autoreactive T cells"

**Supplementary Figure 1**

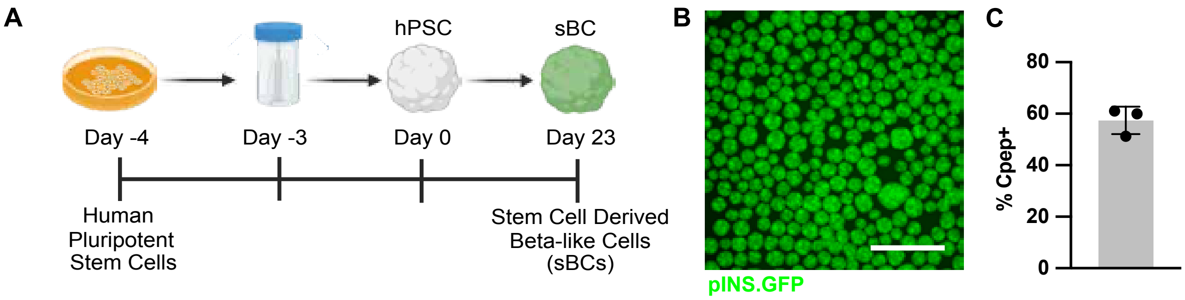


**Supplementary Figure 1: Directed differentiation of stem cell-derived beta cells.** (A) Schematic of the directed differentiation protocol of human pluripotent stem cells towards sBCs in 3D spinner flasks over 23 days. (B) pINS.GFP reporter on day 23 sBCs. Scale bar = 100 $\mu$m**.** (C) Percentage of C-peptide$^{+}$ cells at day 23.

### Supplementary Figure 2

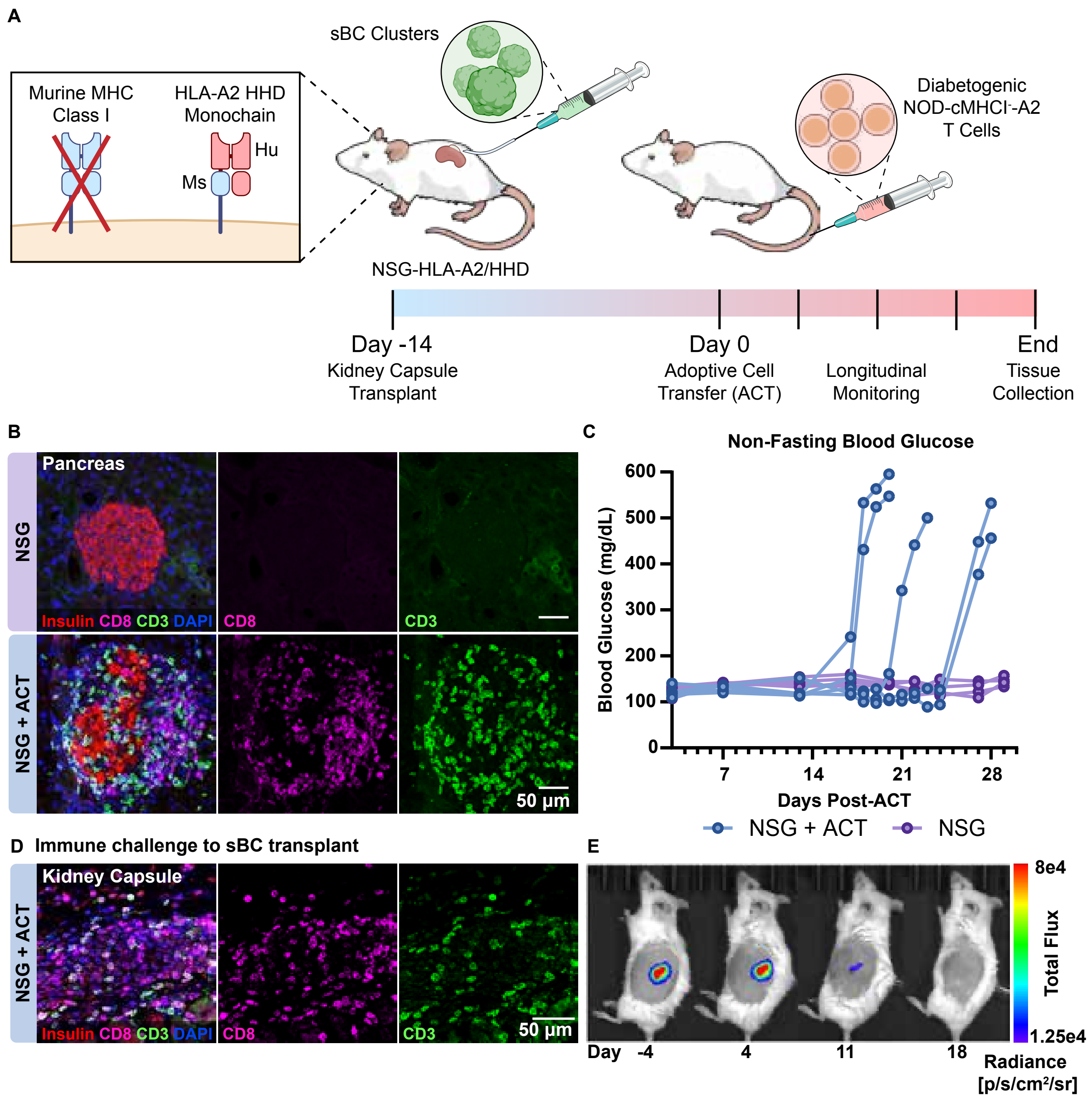


**Supplementary Figure 2: Humanized-HLA Adoptive Transfer Model.** (A) Diagram of mouse model. On day -14 NSG-HLA-A2/HHD are transplanted with human sBC clusters. In these mice, the MHC Class I is knocked out and replaced with a humanized HLA-transgene. After sBCs are engrafted, 5e6 activated $CD3^+$ T cells from a diabetogenic NOD-cMHCI$^-$-A2 mouse are injected i.v. (B) Representative confocal images of pancreas sections from naïve and adoptively transferred mice immunostained for insulin, CD8, and CD3 and counterstained with DAPI. Scale bar is 50 µm. (C) Blood glucose was monitored over 4 weeks, showing conversion to diabetes following adoptive cell transfer. (D) Kidney capsule section showing trafficking of $CD8^+$ T cells to luciferase expressing sBC grafts without microgels, and graft destruction as monitored by (E) IVIS over the course of the study.
